# Agent-Based Modeling Reveals Possible Mechanisms for Observed Aggregation Cell Behaviors in *Myxococcus xanthus*

**DOI:** 10.1101/320291

**Authors:** Zhaoyang Zhang, Oleg A. Igoshin, Christopher R. Cotter, Lawrence J. Shimkets

## Abstract

*Myxococcus xanthus* is a soil bacterium that serves as a model system for biological self-organization. Cells form distinct, dynamic patterns depending on environmental conditions. An agent-based model (ABM) was used to understand how *M. xanthus* cells aggregate into multicellular mounds in response to starvation. In this model, each cell is modeled as an agent, represented by a point-particle and characterized by its position and moving direction. At low agent density, the model recapitulates the dynamic patterns observed by experiments and a previous biophysical model. To study aggregation at high cell density, we extended the model based on the recent experimental observation that cells exhibit biased movement towards aggregates. We tested two possible mechanisms for this biased movement and demonstrate that a chemotaxis model with adaptation can reproduce the observed experimental results leading to the formation of stable aggregates. Furthermore, our model reproduces the experimentally observed patterns of cell alignment around aggregates.

**Author summary:** Collective self-organization of cells into multicellular structures is important for lifestyle of many bacterial species. *Myxococcus xanthus* bacterium is a model system for studying this self-organization. In this work, we investigate how in response to starvation *M. xanthus* cells aggregate into multicellular mounds. A recent study identified the key cellular behaviors that are necessarily for the aggregation but the mechanisms of these behaviors remained unclear. To uncover these mechanisms, we developed a computational model that simulates interactions among a large number of cells. The results demonstrate that the observed bias in the cell reversal times as they move towards the aggregates can be explained by chemotaxis model. In this model cells secrete a chemical signal and respond to it via a partially-adapting biochemical network. The resulting aggregation dynamics are in good agreement with the experiments. Furthermore, chemotaxis signaling model reproduces the experimentally observed patterns of cell alignment around aggregates. On the other hand, an alternative model, based on contact-dependent signaling between cells, fails to aid in aggregation. Thus our models make important predictions about the cellular interactions that drives multicellular aggregation and can serve as a basis to investigate a wider range of developmental mutant strains.

## Introduction

Multicellular self-organization is widely studied due to its biological significance across all kingdoms of life [1–4]. For example, the dynamic organization of biofilms formed by the Gram-negative bacterium *Myxococcus xanthus* depends on the ability of these cells to sense, integrate, and respond to a variety of intercellular and environmental cues that coordinate motility [5–12]. Under nutritional stress, *M. xanthus* initiates a developmental program that stimulates cells to aggregate into multicellular mounds that later fill with spores to become fruiting bodies [13,14]. Despite decades of research, the mechanistic basis of this aggregation behavior in *M. xanthus* is not fully understood.

*M. xanthus* is a rod-shaped bacterium that moves along its long axis with periodic reversals of direction [15]. When moving in groups, cells align parallel to one another due to steric interactions among cells and their ability to secrete and follow trails [13]. Notably, mutations that abolish direction reversals affect collective motility and alignment patterns [16]. Coordination of cellular reversals and collective cell alignment are crucial for multicellular self-organization behaviors [17–19].

Mathematical and computational modeling efforts have long complemented the experimental studies to test various hypotheses about how aggregation occurs [20–24]. However, most modeling research has focused on the formation of large, terminal aggregates rather than the dynamics of aggregation. Furthermore, they have been aimed at elucidating a single, dominant mechanism that drives aggregation. In contrast, our recent work employed a combination of fluorescence microscopy and data-driven modeling to uncover behaviors that drive self-organization [1]. These mechanisms were quantified as correlations between the coarse-grained behaviors of individual cells and the dynamics of the population [1]. Thereafter, parameterless, data-driven models demonstrated that the following observed behaviors are necessary to match the observed aggregation dynamics: decreased cell motility inside the aggregates, a biased walk toward aggregate centroids, and alignment among neighboring cells and in a radial direction to the nearest aggregate [1]. Despite the success of these approaches, the mechanistic bases of these behaviors remain unclear. For example, it is not clear how cells could detect the aggregates to align in their radial direction and extend their reversal time when moving towards the aggregates.

*M. xanthus* produces both contact-dependent signals and chemoattractants. An example of a contact-dependent stimulus is the stimulation of pilus retraction upon the interaction of a pilus on the surface of one cell with polysaccharide on the surface of another cell. This interaction is required for one of the two motility systems deployed by *M. xanthus* [25]. Endogenous chemoattractants are also produced and are known to cause a biased walk similar to that observed during aggregate development [6,26]. The chemoattractants may be lipids, since *M. xanthus* has a chemosensory system that allows directed movement towards phosphatidylethanolamine and diacylglycerol [27].

Here we develop a mechanistic agent-based model (ABM) that aims to test possible mechanisms for the observed cell behaviors. In particular, we examine whether contact-based signaling or chemotaxis can explain the longer reversal times for cells moving toward the aggregates as compared to cells moving away from the aggregates. Furthermore, we explore whether a previously developed cell-alignment model [13] can be scaled up to the proper cell density during aggregation and whether mechanisms postulated in that model – specifically, local alignment and trail-following – are sufficient to explain observed patterns of cell alignment.

## Results

### A phenomenological model matches the patterns of cell alignment

Our recently developed collective-alignment model is based on a biophysically realistic model of flexible *M. xanthus* cells that replicates aspects important for cell-cell interactions, including excluded-volume physics that prohibit overlap between cells in physical space and trail-following behavior [13,14]. This model can explain experimentally observed alignment patterns for wild-type cells and non-reversing mutants at low cell density (0.08 cell/μm^2^, i.e packing fraction is about 25%) [16]. However, the model formalism is too complex to efficiently simulate large numbers of cells. Furthermore, while this model does not allow overlap between cells, at high cell densities the overlap is unavoidable. Therefore, as a first step we developed a phenomenological model of cell alignment. The model details are given in Methods and briefly summarized below.

In our ABM, each agent represents a cell as a self-propelled particle on a 2-D surface with a center position of (*x*(*t*), *y*(*t*)) and orientation *-π*<θ<*π*. The simulations are conducted on a rectangular 2-D area with periodic boundary conditions. At each time-step, agents move in the direction of their orientation and turn to align with their neighbors and any trails in the area. No hard-core, excluded-volume repulsion between agents is explicitly modeled (agents are essentially zero volume). However, to avoid biologically unrealistic agent densities, repulsion between agents in a direction perpendicular to their long axis is introduced. To model alignment with neighbors, we used a phenomenological approach based on Sliusarenko et al. [28]: each agent changes its orientation to align with their neighboring cells at every time step. To model alignment with trails, we used the same approach as in Balagam et al. [13], in which cells turn their long axes to nearby areas with a maximum density of a trail matrix.

To compare the phenomenological model with a detailed biophysical model [13], we first simulated reversing and non-reversing agents at low density (0.08 agent/μm^2^, i.e. ~25% packing fraction) with our ABM in a 400 μm × 400 μm domain with periodic boundary conditions and other parameters set according to Ref. [13]. We found that our model captures the differences in patterns of aligned agent groups (clusters) between reversing and non-reversing agents (Fig. 1). For non-reversing agents, the previous biophysical model [13] shows that agents form isolated clusters (Fig. 1A). Our model shows a similar pattern (Fig. 1B). For reversing agents, the previous model [13] shows that agents form an interconnected, mesh-like structure (Fig. 1C). Our model also shows this pattern for reversing cells (Fig. 1D). These clustering patterns are also observed in experiments for reversing and non-reversing cells [16]. To quantitatively compare the results of the new model with the previous biophysical model, we computed the average number of agents in a cluster as a function of overall agent density (Fig. S1). Our model gives similar results compared with the previous biophysical model -- as expected, higher cell density leads to more agents in a cluster. Therefore, our model quantitatively matches the detailed biophysical model, but is computationally efficient enough to simulate very high cell densities.

**Fig 1:**
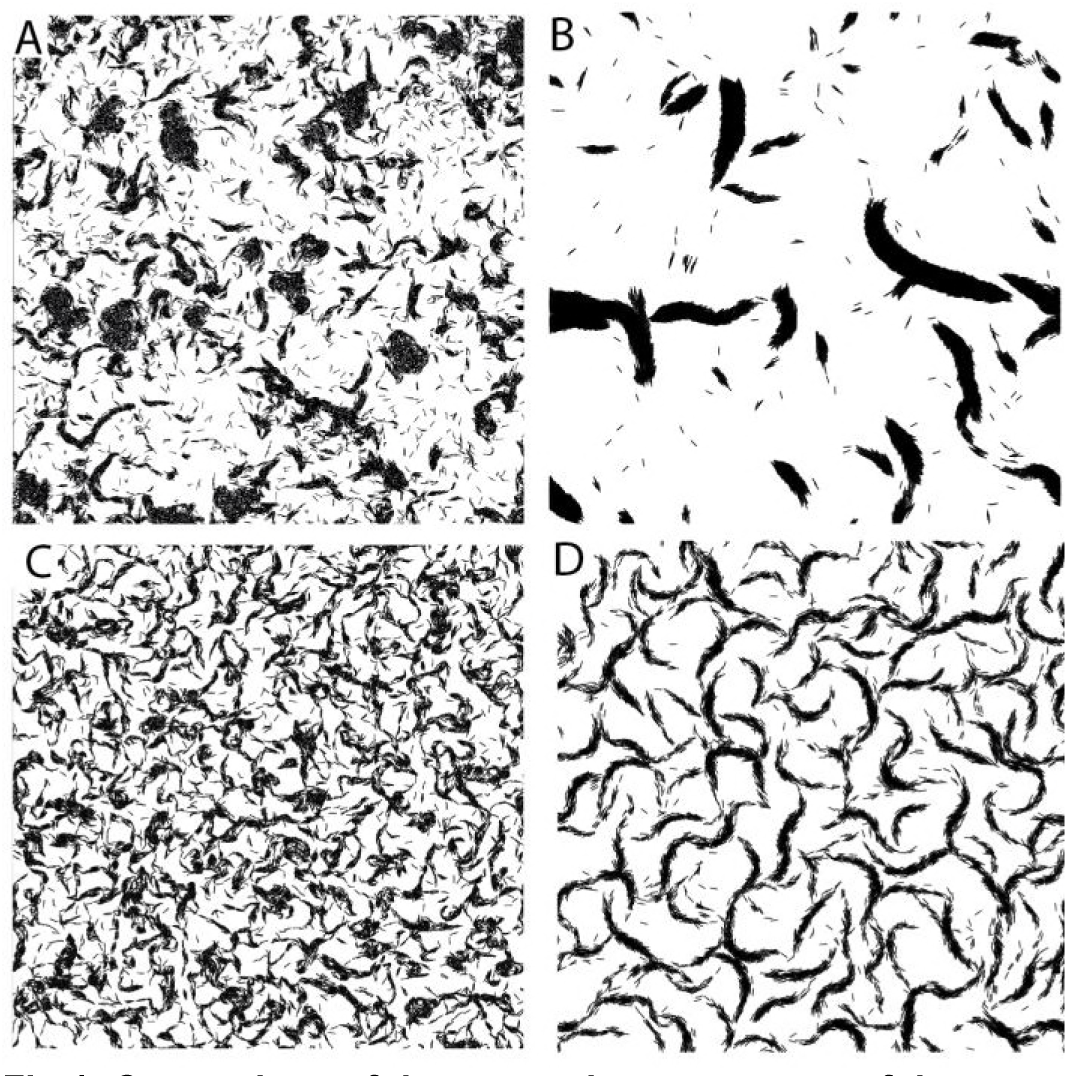
Comparison of the agent cluster patterns of the Balagam et al. [13] biophysical model with our model. The area shown is 400μm × 400μm with the agent density of 0.08 agent/μm^2^. (A,B) Simulation of non-reversing agents of the Balagam et al. biophysical model(A) and of our new model (B). (C,D) Simulation of reversing agents of the Balagam et al. model (C) and of our new model (D).

### Density-dependent motility decrease and cell alignment are not sufficient for aggregate formation

Several previously published models of aggregation were based on the hypothesized “traffic-jam mechanism”, in which cells slowdown in regions of high density [28,29]. This slowdown further increases local cell density leading to a positive feedback loop that drives cell aggregation. Cell tracking experiments by Sliusarenko et al. reported an ~80% reduction of cell speeds in high cell density areas and used these measurements in the model to reproduce aggregation patterns [28]. However, subsequent comparison of aggregation dynamics between the traffic-jam model and experimental results revealed that a traffic jam alone is insufficient to drive aggregation. While the model can produce aggregates of similar size compared with experiments [29] it fails to display the observed [30] dynamic properties of aggregates, such as moving, merging and splitting, nor does it reproduce the correct time dynamics and extent of aggregation. To investigate how different mechanisms of cell alignment would affect the aggregation dynamics, we performed multiple simulations of the traffic-jam model. Following up on the results of [28], agents reduce their speed by 80% if the local cell density is above 0.6 agent/μm^2^. Initially, agents are randomly distributed with density 0.3 agent/μm^2^.

In the first model we consider no local alignment; all the agents choose random orientations. Running the simulations of this model for 10 hours, the natural time-scale of aggregation in Ref. [1], we observed the emergence of very small aggregates (Fig. 2A). If we continue the simulations for longer times, the aggregation process becomes unrealistically long: even after 50 hours, some aggregates are still growing (Fig. S2A).

**Fig 2:**
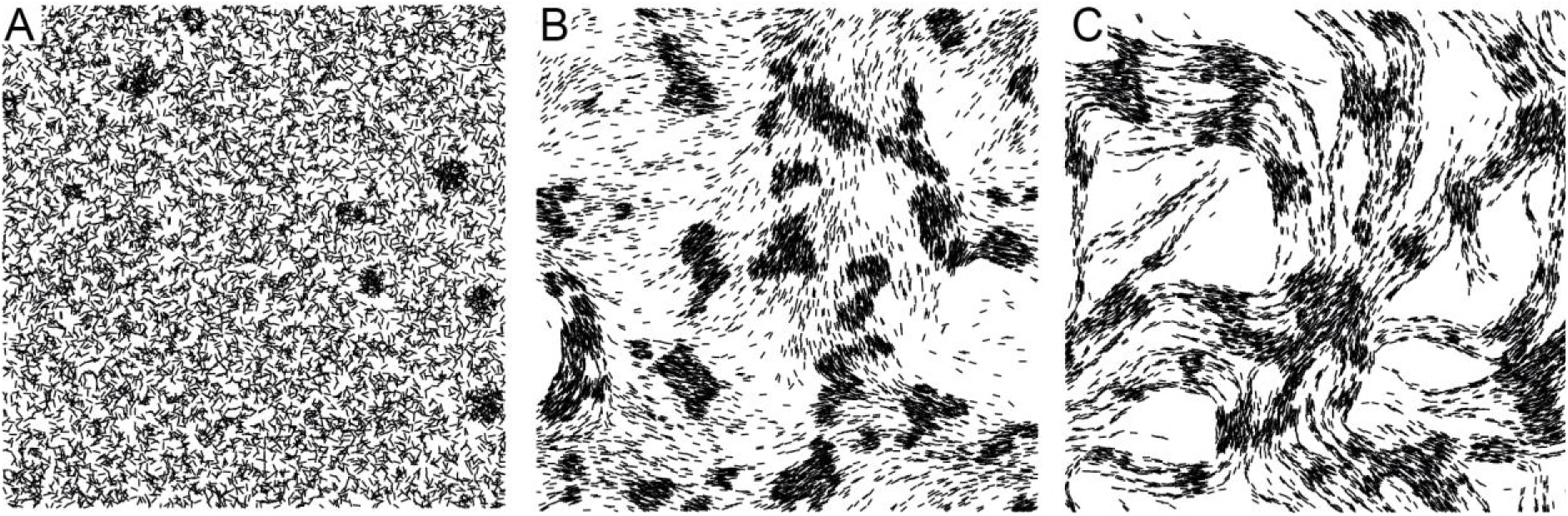
Comparison of the collective behaviors of different models. (A) Traffic jam model. Simulation result at 10 hours. (B) Traffic jam model with local alignment. Simulation result at 10 hours. (C) Traffic jam model with local alignment and slime trail following. Simulation result at 10 hours. All simulations have agent density 0.3 agent/μm^2^ and 400μm × 400μm area. To make aggregates more visible, only randomly selected 20% of agents are plotted.

In the second model we also consider local alignment: agents now actively align their long axis with nearby agents. There are more aggregates in Fig. 2B and their size is larger than the observed results in Fig. 2A. Therefore, in agreement with the results of Ref. [2,28], local alignment helps with aggregation. However, examining the patterns of aggregation dynamics (Fig. S2B) we note that aggregation is still slower than experimentally observed [1]. Moreover, most of the aggregates do not have the circular shape observed in experiments [1,30] and aggregates do not move, merge, or split.

The results of Ref. [3,13] indicated that trail following further aids aggregation. To test the effect of trail-following on the aggregation, we include these effects into our second model. As shown Fig. 2C, by 10 hours cells converge into streams and form small aggregates at the intersections of streams. However, these aggregates are unusually small and do not grow in size given more simulation time (Fig. S2C).

From these results we conclude that a traffic-jam model is not sufficient to form aggregates even when proper alignment mechanisms are included in the model. Moreover, recent results from Cotter et al. showed that cell motility reduction at high cell densities is only about 60% [1] further impeding the traffic-jam model. Therefore, we can conclude that the motility reduction is insufficient to drive aggregation. This agrees with the conclusion of Cotter et al. [1] who demonstrated that biased walk toward the aggregate center is essential to match experimentally observed aggregation dynamics. However, the biochemical mechanism of the bias has not been identified. In what follows, we test two alternative mechanisms for this biased walk.

### Contact signal based model does not lead to stable aggregation

Many previous studies suggest that the reversal frequency is affected by contact with other cells [4,5,18,31,32]. For example, Mauriello et al demonstrated that when *M. xanthus* cells make transient side-to-side contact, they exhibit increased cellular reversals [4]. Earlier studies also postulated control of reversals during *M. xanthus* development by cell contact due to C-signal exchange [5,18,31,32]. Moreover, mathematical models based on the contact-dependent reversal induction successfully reproduce traveling wave patterns formed by *M. xanthus* cells during development [33]. Despite all the work postulating reversal control via C-signal exchange, no molecular mechanism has been identified.

We begin by testing the hypothesis that *M. xanthus* cells employ a contact-based mechanism to perform a biased walk. To test this hypothesis using our ABM, we assume that an agent’s run duration can be affected by nearby agents’ run directions. If nearby agents are moving in the opposite direction, the target agent will have a shorter run duration. If nearby agents are moving in the same direction, the target agent will have a longer run duration. In this way, when cells form a stream, it is likely that more cells will run in one direction to produce a net flow of cells. When the streams of cells intersect, an aggregate can form at the intersection.

To decide how reversal period is influenced by the direction of nearby cells, we follow the data-driven modeling approaches of Cotter et al [1]. For simplicity, we first run a simple 1D, open-loop simulation. In this simulation, we assume there is an aggregate in the middle of the simulation domain. This model is completely data-driven; agents choose their behaviors from an experimental dataset from Cotter et al [1]. As in Ref. [1], agent behavior is sampled from the recorded dataset of cell behaviors conditional on how far the agent is from the aggregate and the agent’s direction relative to the aggregate. As expected, the biased walk led to agents accumulating in the region of the postulated aggregate. As a result, the density of agents moving toward the aggregates is slightly higher than the density of agents moving away from the aggregates (Fig. 3A up right). With this result, we asked whether the run bias can result from contact-based signaling that relies on an agent sensing nearby agents that move in same (*n*_+_) or opposite directions (*n*_−_).

**Fig 3:**
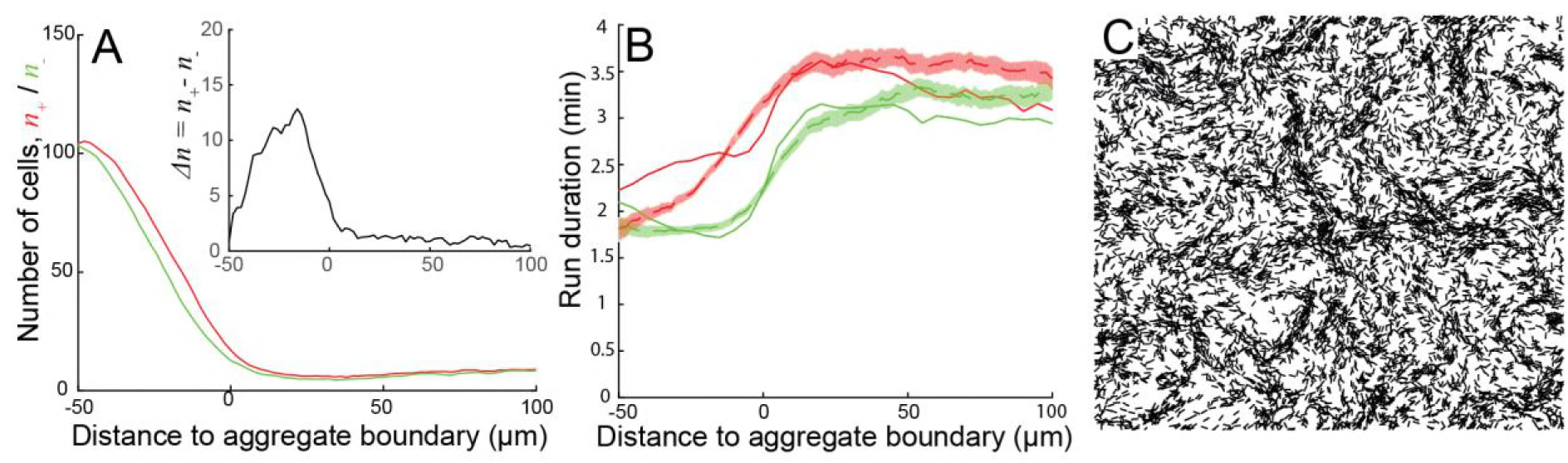
Results of the contact signal model simulations. (A) Agent density (i.e. number of neighbor agents within 3μm of each agent) around the aggregate in 1D open-loop simulation as a function of distance to aggregate boundary. The number of agents that move towards the aggregate center (n_+_) shown in red whereas the number of agents moving away from the aggregate center (n_−_) is in green. Insert shows the difference between the two Δn = n_+_ − n_−_. Results are averaged over 23 hours time. (B) Experimental run duration data fitted to the proposed contact-signaling model. Red lines indicate run durations of cell moving toward aggregates. Green lines indicate run durations of cells moving away from aggregates. Solid lines are fitted run durations and dashed lines are experimental results. Red and green shaded areas are bootstrapped 95% confidence intervals calculated from experimental results. Experimental data is from [1]. (C) Closed-loop simulation of ABM with contact-dependent signal at 10 hours. Run duration of agents is computed using the fitted parameters from (B). Initially, agents are randomly distributed across the simulation domain with a density of 0.3 agent/μm^2^. As in Fig 2, simulation area is 400μm × 400μm and only 20% of agents are plotted.

Given the relatively small difference between *n*_+_ and *n*_−_ we argue that cell’s reversal period must be sensitive to the sign and the magnitude of the difference in the two. Therefore, we define the signal Δ*n* = *n*_+_ − *n*_−_. In addition to this signal, the reversal period must also depend on the total cell density *ρ_c_* = *n*_+_ + *n*_−_ to ensure a shorter reversal period inside the aggregates regardless of the direction. Therefore, we look for a fit to bias data using

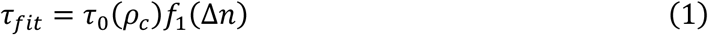

Here the first term will be a decreasing function of *ρ_c_*, e.g.

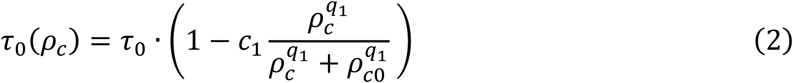
where *τ*_0_ is the mean reversal period at low cell density, *c*_1_,*q*_1_,*ρ_c_*_0_ are parameters chosen to fit the experimental data showing more frequent reversals inside the aggregates where cell density is high. The factor *f*_1_ is responsible in the bias in runs, i.e. the difference in reversal periods for cells going towards or away from the aggregates. We use

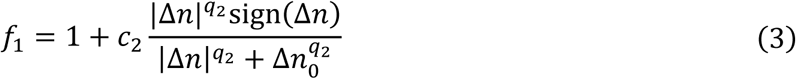
where *c*_2_,*q*_2_,Δ*n*_0_ are parameters chosen to provide the best fit to experimental data. Using Eq. (1)–(3) we can fit the reversal period bias (Fig. 3B). From the results we can conclude that within the framework of this open-loop, 1D model, this contact signal mechanism can fit the run duration data acquired from experiments. However, it is not clear if this mechanism will work in the more rigorous 2D, closed-loop model.

To investigate whether contact-dependent reversal period modulation will aid aggregation, we performed 2D simulations using the model with the directional sensing mechanism. In this model, each agent’s reversal period is given by Eq. (1–3). In Fig. 3C we see that after 10 hours of simulation, this model does not produce any aggregates. Moreover, no stable aggregates form even at longer simulation times (Fig. S3) Therefore, we conclude that the contact-based signal mechanism does not reinforce aggregation and an alternative mechanism for the biased walk of *M. xanthus* cells is needed.

### A chemotaxis model produces biased movement similar to that observed in experiments and helps with aggregation

Previous experiments have showed that *M. xanthus* cells can move up a gradient of lipid extracted from starving cells by altering their reversal period and adapting to the chemical signal [6]. This process is dependent on chemosensory systems similar to that found in other bacteria [7]. We propose that the movement bias during aggregation is caused by chemotaxis and adaptation towards a chemical signal produced by cells. In our model, a large number of cells in an aggregate produce a concentration gradient of a chemical signal that diffuses away. Cells outside the aggregate are then attracted to the aggregate though a biased walk typical of bacterial chemotaxis systems [34].

Based on previous works on chemotaxis adaptation [8–10], we developed our phenomenological model for the adaptation network for chemotaxis based on the integral feedback control (Fig. 4A, see *Methods* for details). We let agents produce and detect signaling molecule (concentration denoted as *ρ* in Fig. 4A) which diffuses and decays with time. At the high cell densities inside aggregates, a gradient of *ρ* surrounding the aggregate can bias cell movement toward aggregation centers. Signal *ρ* excites an internal signal of the agents *y*_1_, which corresponds to the activation of receptors in cells. *y*_2_ is the integral of *y*_1_, and feeds back to the system to inhibit *y*_1_ and deactivate the chemoreceptors (е.g. via demethylation). Therefore *y*_1_ and *y*_2_ form an integral feedback loop for adaptation. Next, we let the reversal period depend on *y*_1_ and *y*_2_:

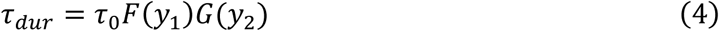

**Fig 4:**
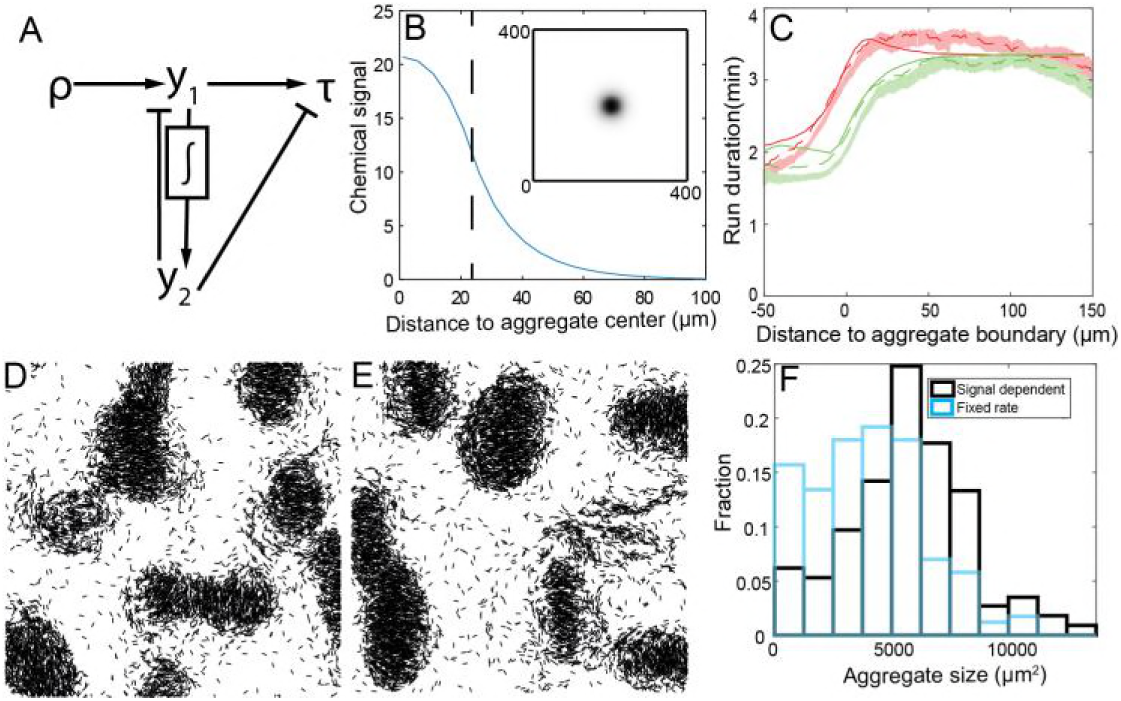
Results of a chemotaxis model. (A) The adaptation network used in our chemotaxis model. (B) An example of a chemical density distribution. The chemical is produced in a circle in the center with fixed decay and diffusion rates. Upper right insert: chemical density in the simulation region at steady state. Main figure: chemical density as a function of distance to aggregate center at steady state. The dashed line marks the boundary of the assumed aggregate. (C) Comparison of run duration biases produced by simulation (solid lines) and experimental data (dashed lines). Red lines are runs pointing toward the aggregate centroid, green lines are runs pointed away from the aggregate centroid. Shaded areas are bootstrapped 95% confidence intervals of experiment data. (D) Simulation of chemotaxis model. Run duration is acquired from fitted duration in (C). Initially, agents are randomly distributed across the simulation domain with density 0.3 agent/μm^2^. Agents are run for 5 hours without chemotaxis to become fully aligned. Picture is taken after 10 hours of simulation with chemotaxis. Simulation area is 400μm × 400μm and only 20% of agents are plotted.

Where *F*,*G*: **R**→(0,∞). Based on the experimental data [1], agents have shorter run duration inside aggregates. Since inside aggregates the signal *ρ* is higher, the steady state of *y*_2_ is also bigger for agents inside aggregates than agents outside aggregates. Therefore, *G* negatively depends on *y*_2_. For *F*(*y*_1_), when agents move up the chemical gradient, *ρ* is increasing, the excitation from *ρ* is stronger than the inhibition, *y*_1_ > 0. and similarly, when agents move down the chemical gradient aggregates, *y*_1_ < 0. Therefore, to make agents bias to longer reversal times as they move up the chemical gradient, *F* needs to be positively dependent on *y*_1_. Considering *F*, *G* > 0 for all possible inputs, we chose

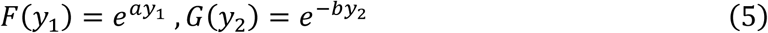
where *a*,*b* are chosen to fit the experiment data.

The gradient profile around the aggregate is determined by the diffusion and degradation of the signaling molecule. We the diffusion coefficient of lipids as a diffusion coefficient [35] and fit the value of the degradation constant to match the range of bias around the aggregates. To determine values for the parameters in our model that match the bias observed in Ref. [1], we performed several 2D simulations. In each simulation, the chemical signal is produced in a circle at the center of the simulation domain assumed to be an aggregate. The sizes of the circle are different in each simulation and are determined by the distribution of aggregates measured in experiments [1]. Fig. 4B shows a chemical profile produced in one of these simulations. To measure the biased walk of agents, we average the run durations based on run direction and distance to aggregate boundary for all simulations. The results in Fig. 4C show that this mechanism produces a bias that matches experimental observations.

To investigate whether the chemotaxis model aids aggregation, we performed 2D simulations using the chemotaxis model. The simulation was started 5 hour run to allow agents time to fully align before the onset of aggregation. During this period, agents do not produce the chemical signal or perform biased movement, they only align with their neighbors and trails. After 5 hours, agents start to produce the chemical signal and perform biased movement. After 10 hours of simulation with chemotaxis, agents form aggregates with clear boundaries (Fig. 4D). However, we noted that the average size of the aggregates in such simulations (mean area ≈ 4000 μm^2^) are somewhat smaller than those in the experiment (mean area ≈ 6000 μm^2^). Moreover, the run durations computed from closed-loop simulations did not quite match the data on Fig. 4C (data not shown). We argued that to correct this issue we need to increase agent’s bias toward larger aggregates, and that can be achieved if we introduce a positive feedback in the chemical signal production: agents produce more chemical signal when in the environment of with high signal concentration (see Methods for detailed implementation). We find that such positive feedback in signal production can produce larger aggregates (Fig. 4EF) and further matches the experimentally observed trends the runs bias (Fig. S7). Thus, we conclude that this chemotaxis model can produce biased movement and stable aggregates. Therefore, we’ve decided to perform larger scale simulation and quantitatively compare the results of the model with these of Cotter et al. [1]

### A chemotaxis model qualitatively reproduces cell alignment patterns

One of the most unexpected findings of Cotter et al. [1] was a finding that *M. xanthus* cells align their moving direction in a radial direction around the nearest aggregate and this alignment aids aggregation. On the other hand, cells inside aggregates and near the aggregate boundary align in the tangential direction, i.e. rotate around the aggregate, whereas cells further away from the aggregate become aligned with the aggregate [1]. To see if our ABM with alignment, trail following, and chemotaxis can reproduce this alignment pattern, we performed simulations using the chemotaxis model as described in last section. Since the microscope field of view in experiment is 986 μm × 740 μm, we increased our simulation domain to 1 mm × 1 mm to better compare with experiment.

First, we compared the alignment patterns of our model and experiment (Fig. 5A-D) Alignment of cells to aggregate centroids is quantified by measuring the alignment of run vectors (in direction *θ_run_*) and vectors pointing toward the nearest aggregate centroid (in direction *θ_agg_*) using cos(2(*θ_run_* − *θ_agg_*)) [1]. Negative alignment to aggregates indicates that cells are rotating around the aggregate; positive alignment to aggregates indicates that cells are aligned toward the aggregate. Our results (Fig. 5A) show that, inside the aggregates the alignment is negative while further away from the center, agents are aligned toward the aggregates as observed experimentally [1]. This result shows that our model is sufficient to explain cell alignment within and around aggregates. Furthermore, we measured alignment of agents to aggregate centroids at different times in the simulation (Fig. 5B). We observed that the average alignment to aggregate centroids is decreasing with time. We think this is because the cell streams pointing towards aggregate centroids are moving into aggregates. Therefore, as time increases, there are fewer cell streams outside of aggregates pointing toward aggregate center. Besides, as the aggregates grows, there are more cells near the boundary circulating around the center. Therefore, the overall alignment to center is becoming more negative as time increases. We test this prediction by reprocessing the data from Cotter et al. [1] within different time-windows. The results demonstrate the same trend as predicted in the model (Fig. 5C). Fig. 5D shows the local alignment of run vectors in experiment and simulation. In our simulations, the local alignment of cells is slightly weaker than the experiment.

**Fig 5:**
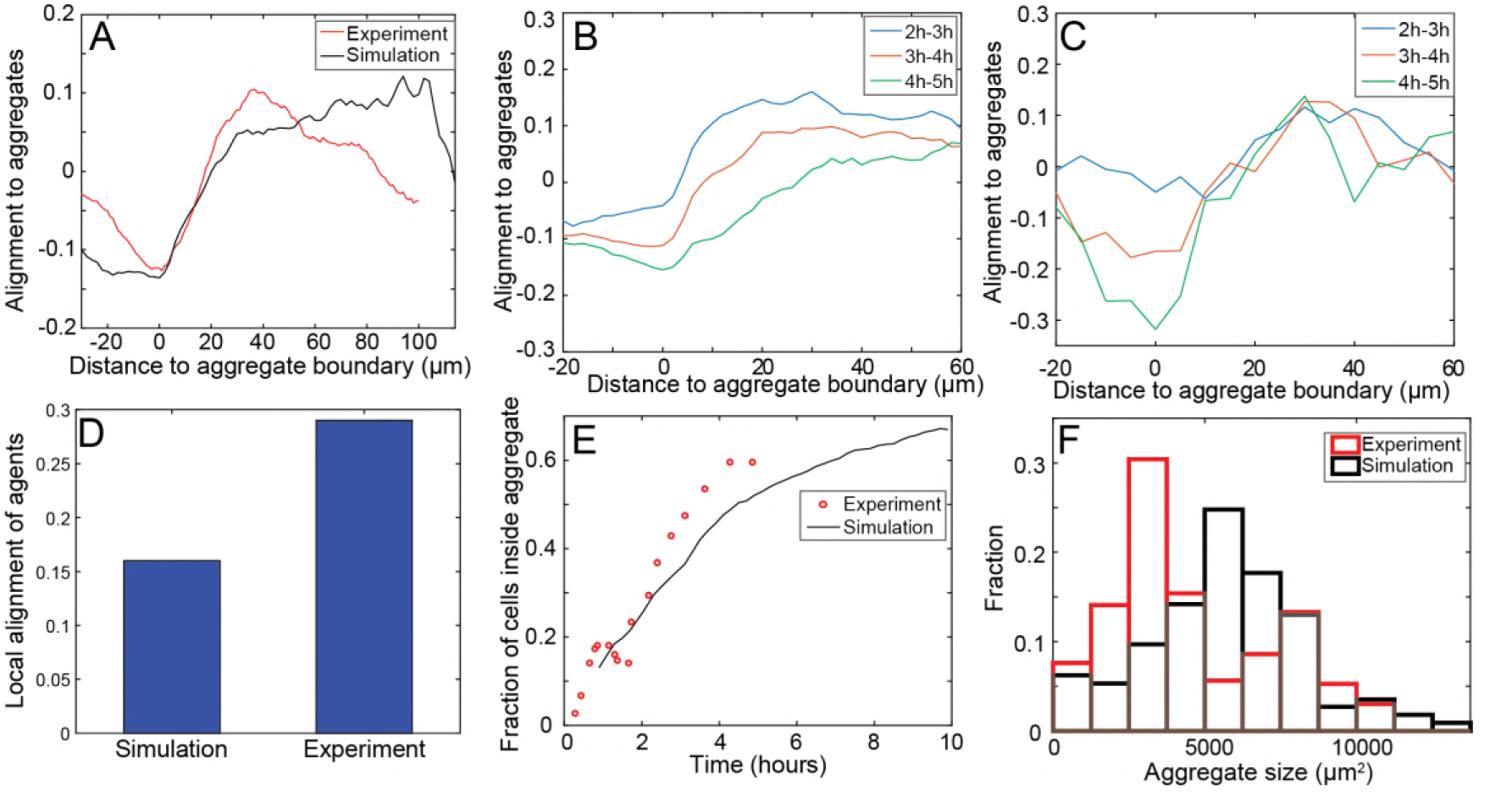
Comparison between simulation and experimental results. Simulation area is 1mm × 1mm. Simulation is prerun for 5 hours without chemical production to align agents. (A) Alignment strength of run vectors with vector pointing toward nearest aggregate centroid. Red line is experimental result and black line is simulation results. Negative distances indicate that the run began inside an aggregate. Values may span (−1, 1) where 1 indicates vectors are parallel. Likewise, −1 indicates vectors are perpendicular. (B) Alignment strength of run vectors with vector pointing toward nearest aggregate centroid at different times in simulation. Results are averaged for run vectors within 1 hour time window. (C) Alignment strength of run vectors with vector pointing toward nearest aggregate centroid at different times in experiment. Results are averaged for run vectors within the 1 h time window. (D) Run vector alignment strength with neighboring run vectors that occurred within ±5 min and 15 μm of different run durations. Values may span (−1, 1) as in (A). (E) Fraction of cells in aggregate in experiment (red) and simulation (black). (F) Aggregate size distribution at 5 hours in experiment (red) and at 8 hours in simulation (black). The measurement time point in simulation is chosen when the fraction of cells inside aggregate reaches 60%. The aggregate size distribution is measured for multiple simulations with more than 100 aggregates.

Comparison of aggregation dynamics in our model to the experimental observations in the Ref. [1] indicate similar aggregation rates. (Fig. 5E). We note that the quantitative agreement between the model and the experiments can be further improved by the offset between the time-point of 0 hours between the two; we assumed that in the model t=0 hours is the time when agents start chemical signal production. In simulation, it takes about 8 hours for the fraction of cells inside aggregates to reach about 60% as compared to 5 hours in the experiments. Over 10 hours the model gets over 65% of the cells into the aggregates. Distribution of aggregate sizes (Fig. 5F) further demonstrates quantitative agreement between the simulations (black bars) and experiments of Ref. [1] (red bars). However, the agreement is not perfect as the model produces slightly more large aggregates (over 5000 μm^2^) and fewer small aggregates (below 5000 μm^2^). This agreement can perhaps be made better by further tuning the positive feedback in the chemical signal production but is beyond the scope of this work. Nevertheless, developed model shows remarkable agreement with most of the features of aggregation dynamics quantified by Cotter et al. [1]

## Discussion

In this paper, we developed a mechanistic ABM that matches many dynamic features observed during aggregation and provides plausible mechanistic explanations for the cell behaviors uncovered in our previous work [1]. Given the importance of cell alignment in aggregation, we started by developing a phenomenological framework that approximates interactions of the previous biophysical model [13]. The framework can reproduce the different cell-alignment behaviors of reversing and non-reversing *M. xanthus* cells at low cell densities [16] and can be scaled to much higher aggregation cell densities with multiple cell layers. Next we explored possible mechanisms of the biased walk toward the aggregates [1]. We showed that a chemotaxis model but not a contact-based signal model produces stable aggregates and the biased movement in our model. Remarkably, the resulting model also matches the experimentally observed cell alignment patterns.

Why would chemotaxis but not contact-signaling model reproduce aggregation patterns? We argue, that the failure of a contact-based signal model to produce stable aggregates is due to its high sensitivity to noise. As seen in Fig. 3A, the signal used in the model – difference between the densities of cells going towards vs. away from the aggregates is much smaller in comparison with the local cell density. Therefore, the fluctuations in local cell density could significantly affect the input signal. Further, this signal rapidly changes in time with local cell density. On the other hand, a secreted chemoattractant is directly proportional to cell density and changes gradually over time. The ability of cells to detect both the value and time derivative (via adaptive network) of the signal allows cells time to aggregate in high chemical signal areas.

Notably, our simulations match the observations from Cotter et al. [1] that away from the aggregate boundaries, cells tend to align radially to aggregate centers (Fig. 5A) while closer to the aggregate boundaries tangential (circumferential) orientation patterns are observed (Fig. 5A). We note that in the model the orientation of agents is not directly affected by aggregate positions or agent density or chemical signal gradients. Therefore, our model predicts that the observed alignment patterns are results of the two major factors affecting collective cellular alignment dynamics: alignment to nearby cells and to the secreted trails. As a result of these factors, cells align and form streams prior to the initiation of aggregation. Once aggregation initiates, cells predominately move along the streams and, as a result, aggregates tend to form in the regions where these streams intersect. Since cells tend to move toward aggregates, aggregate size and cell density inside aggregates increases. At the high cell densities that exist inside an aggregate, cells may not be able to move any closer to the aggregate center (due to local repulsion), instead start moving in the circumferential direction. Further away from the aggregates, cells continue to move along the streams toward or away from the aggregate, which leads to radial orientation patterns with respect to aggregates. The radial orientation patters become weaker as the density of cells outside the aggregate decreases.

At this point, we focused mainly on qualitative agreement between our model and experimental aggregation patterns but nevertheless got quantitative agreement of multiple aggregate properties. However, we argue that this agreement can be further improved in the future work since our models ignore certain aspects of cell behaviors observed by Cotter et al. [1]. For example, the amount of stochasticity in the individual cell behaviors will undoubtedly affect the aggregation dynamics. The results of Cotter et al. showed wide variation in the persistent run durations of cells [1]. In our simulations, we set the noise to be somewhat smaller and did not investigate its effect on aggregation. Investigating the effect of noise on aggregation will be an interesting direction for future work. Furthermore, during our aggregation simulations we assumed time-independent cell behavior but the results of Cotter et al. showed dynamical changes in cell speed and run durations. The effect of this variability will also be systematically investigated in future studies. Notably, the cells display longer reversal periods in the beginning of aggregation[1] and long reversal periods combined with steric interactions and trail following have been shown [13] to enable formation of circular aggregates – cell streams that close on themselves. It remains to be seen if these aggregates can serve as nucleation centers of more mature aggregates. Nevertheless, our model makes important predictions about the cellular interactions that drives multicellular aggregation and can serve as a basis to investigate a wider range of developmental mutant strains.

## Materials and Methods: Agent-based model simulation framework

### Model of *M. xanthus* cell

We developed our model based on the well-known Vicsek model [11]. In our model, each cell is represented as a self-propelled particle (agent) on a 2D surface with a center position of 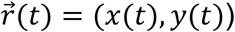, and orientation −π < *θ* < π. Time is updated in the simulation by constant increment Δ*t* = 0.05 min. The simulations are conducted on a rectangular 2D area with periodic boundary conditions. The agent’s position is updated at each time step depending on its velocity and orientation:

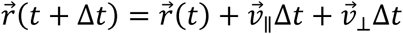

Here 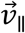 is the cell velocity along its long axis,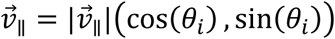, and its magnitude depends on the local cell density *ρ_c_*:

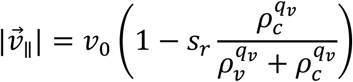

Here *v*_0_ is cell speed at low cell density, *s_r_* is the speed reduction fraction at high cell density, *q_v_* is the power index that controls the curve and *ρ_v_* is the density threshold of speed reduction.

Term 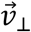 is the cell velocity component perpendicular to cell long axis due to repulsion of other cells. In our model, each agent does not have any excluded volume and therefore and can overlap with other agents. However, in the experiments with low cell density cells do not overlap. To mimic this behavior in our modeling framework, we introduced repulsive interactions between nearby agents. First, we defined perpendicular distance between cell *i* and its neighbor cell *j* as shortest distance between the cell *j* and the line though cell *i* in the direction of (cos(*θ_i_*), sin(*θ_i_*)). It can be computed by projecting the vector connecting two agent positions to 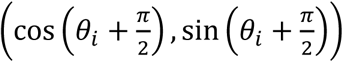:

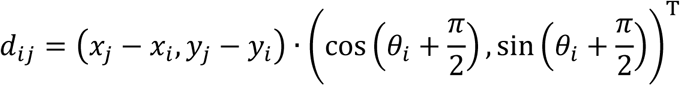

We then use this distance to define a repulsive force that serves to move cells apart from one another and to prevent overlap at lower densities. At higher cell densities, i.e. when cells are in multiple layers, this force will prevent unreasonably high local cell densities. We express this repulsion force 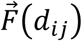 acting on cell *i* as follows:

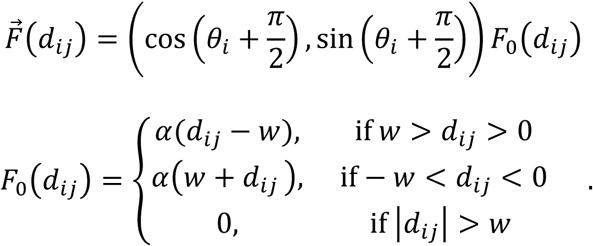

Here *α* is an effective spring constant and *w* maximum interaction range. The latter is set to be 0.5 μm, approximately equal to the cell width.

Every time step, for each cell we compute the perpendicular distance *d_ij_* for all cells *j* near the target cell *i* (within 3 μm radius, which is half-cell length) and then compute the corresponding 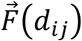. Thereafter, we sum up all the repulsion force of cell *i* and calculate 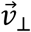 proportional to the force (assuming an overdamped limit, very low Reynolds number):

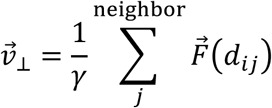

Here, is *γ* viscous drag coefficient. The parameter *ε_rpl_* = *α*/*γ* therefore controls the effectiveness of volume exclusion interaction to push cells apart.

Orientation of the agent (*θ*) changes due to three factors: stochastic fluctuations (noise, stochastic turning), alignment to its neighbors, and cell’s tendency to follow trails:

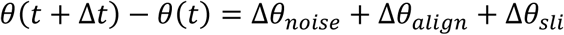

Here Δ*θ_noise_* is the noise term estimated from experimental data [14], Δ*θ_align_* is local cell alignment, and Δ*θ_sli_* is trail-following effect as defined introduced in the following subsections.

### Local cell alignment

In this model, we chose to model nematic (i.e. mod 180 degrees) cell alignment to neighbor cells based on the equations of Sliusarenko et al [28]. The alignment of cell *i* in one time step Δ*t* is calculated as:

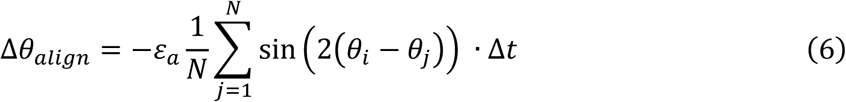

Here *θ_i_* is the orientation of *i^th^* cell, *θ_j_* is the orientation of the *j^th^* cell, which is one of *i^th^* cell’s neighbors. *ε_a_* is the parameter that controls the strength of cell alignment. The summation is done over *N* cells within a 5 μm raidus of *i^th^* cell.

### Trail-following

*M*. xanthus cells tend to follow the trail left by pioneer cells at low cell density [36] and 3D tunnels within the biofilm at high cell density [37]. The exact mechanism for this trail-following by *M. xanthus* cells is currently unknown. As in our previous model [13] we assume that cells actively seek trail-rich regions on the substrate.

We employed the same phenomenological approach as in Balagam et al. [13] to build a trail field (*S*(*x*,*y*,*t*) recording the trail material density at position (*x*,*y*) at time step *t*) covering the entire simulation region. We keep track of its time-evolution using a square lattice grid with grid width equal to the cell width (0.5 μm). During each time step Δ*t* every agent deposit *S_pr_* × Δ*t* trail material at the grid-point closest to its location; *S_pr_* is the trail production rate. Trail material decays exponentially over time with a constant degradation rate *β_m_*, e.g.: *S*(*x*,*y*,*t*+Δ*t*) = *S*(*x*,*y*,*t*) × *e*^−^*^βm^*^∆^*^t^*, *S*. As in Balagam et al. [13], to determine the preferred trail direction (*θ_trl_*): we let each agent detect the trail in a semicircle in front of the agent, *θ_trl_* points to the area with least deviation for agents orientation *θ*(*t*) and with more than 80% of the maximum trail field density. However, within our ABM alignment to this new directions is done with phenomenological approach distinct from Balagam et al. [13] but similar to Eq. (6), i.e. we gradually change the orientation of agent *i* parallel to the preferred direction:

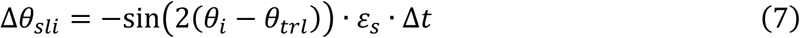

Here, *ε_s_* controls how fast agents align to *θ_trl_*, in other words, it controls the strength of the trail-following effect.

### Periodic reversal of cells

*M. xanthus* cells periodically reverse their travel direction [38]. Following previous work, we let the reversing period length τ obey Gamma distribution [39], the probability density function of reversal period *τ* can be written as:

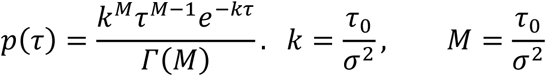

Here, *τ*_0_ is the average period length and *σ* is the standard deviation of period length. To track the time between cell reversals, we introduced an internal timer (*t_c_*) to record how many time steps the cell has been in this reversal period. In simulation, the probability of reversing at *t_c_* = *K* (*K* ∈ *N*) time step is calculated from a probability density function *p*(*t*), written as:

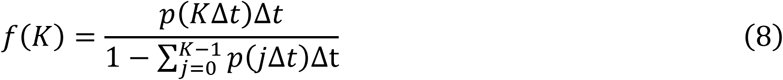

The numerator is the probability of reversing at time step *K*; the denominator is the probability of not reversed in the preceding time step from 0 to (*K* − 1).

### Chemotaxis with an adaptation model

Previous studies with *E. coli* [9] have proposed a mechanism for robust adaptation in simple signal transduction networks. The adaptation property is a consequence of the network’s connectivity and does not require the ‘fine-tuning’ of parameters. Based on this robust adaptation model, Yi et al. [40] gives a standard solution that can achieve perfect adaptation: integral feedback control, in which the time integral of the system error, the difference between the actual output and the desired steady state output, is fed back into the system. This feedback control can be achieved by simple differential equations [40]:

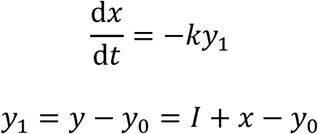

Where *x* is the time integral of system error and *y*_1_ represents the error, which is the difference between the actual output *y* and the steady-state output *y*_0_. *y*_0_ is a constant determined by the enzyme level of the system. *I* is the input signal and *k* is a parameter that determines the adaptation rate, i.e. inverse of adaptation time-scale. In steady state, *y*_1_ = 0, *x* = *y*_0_ − *I*. Therefore, *y*_1_ does not depend on the input signal *I*.

To remove constant *y*_0_ from our model, we set *y*_2_ = *y*_0_ − *x*. Then we have

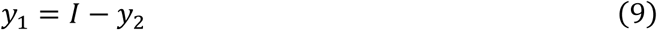

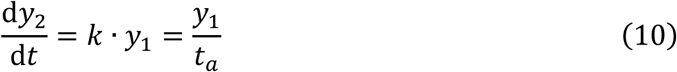

Where *t_a_* is the adaptation time.

For input signal *I*, we let *I* = *f*(*ρ*). Where *f*:[0,∞) → [0,∞) models the first step of signal transduction and *ρ* is the signal in the environment. For function *f*, we employ the same function as in Ref. [8]

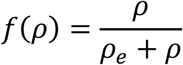

Where *ρ_e_* is a parameter controlling the signal.

Therefore, for any constant signal *ρ* we have the property that 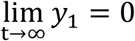 and 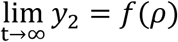.

### Chemical signal diffusion and auto regulated dependent production

In our model, we assume cells produce a chemical signal to aid aggregation. We use a 5 μm × 5 μm square lattice covering the whole simulation region to record the signal. Each cell produces *α*(*ρ*) × Δ*t* amount of signal every time step at its location. Here *α*(*ρ*) is the production rate that is dependent on chemical signal *ρ*. Thus, we have diffusion-reaction equation for the concentration of signal:

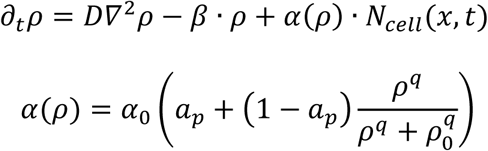

*N_cell_*(*x*,*t*) is number of cells at positon *x* and time *t*, *D* is diffusion coefficient and *β* is decay coefficient. *α*_0_ controls the production rate. *a_p_* ∈ (0,1) controls the dependence of *α*(*ρ*) on *ρ*. We set our diffusion coefficient based on the experiment of lipid diffusion [35]. Moreover, we also performed simulations of different diffusion rates (Fig. S6). In order to keep the chemical gradient the same, we also scaled the decay rate and production rate with the diffusion coefficient. We see that when the diffusion coefficient is too big (3000 μm^2^/min), fewer agents accumulate in aggregates because it is harder to accumulate a chemical signal in the environment. Therefore, the positive feedback is not obvious and the chemical production is insufficient. However, for a diffusion coefficient of 300 μm^2^/min or 30 μm^2^/min, aggregation rates are similar.

### Persistent to non-persistent state transition

Previous work has defined persistent state and nonpersistent state of *M. xanthus* cells [1]: A persistent state was assigned to trajectory segments in which cells were actively moving along their long axis; nonpersistent state was assigned when cell velocity is too small (less than ~1 μm/min) or reversal period too high (greater than ~1 reversal per minute). The probability of cells transiting from persistent state to non-persistent state was measured as a function of distance to aggregate [1]. In our model, we let the probability directly depend on chemical signal *ρ*. We use the following equation to calculate this probability.

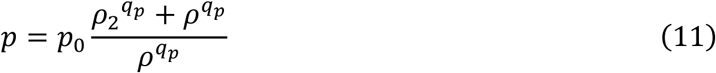

*ρ* is the signal level, *p*_0_, *ρ*_2_, *q_p_* are parameters chosen to fit experiment data. Here we did not differentiate cells’ moving direction for two reasons: 1) the difference in transition probability is small between cells going in and out of aggregates [1], 2) the difference in transition probability does not have a big impact on cells’ motility bias.

We performed simulations on a 400 μm × 400 μm area to fit eq. (11), in order to mimic the signal produced by an aggregate placed in the center. We let the signal be produced in a circle that is located in the center of the simulation domain, with 50 μm radius. The signal also diffuses and decays with time. We used the steady state of signal level to fit eq. (11). Because the signal *ρ* is centrosymmetric, we fit eq. (11) in 1 dimension at *y* = 200 μm.

For nonpersistent state duration, we did not differentiate the moving directions of cells. because in this state cells have very low motility. We think the direction in nonpersistent run is not important.

In the model, we let the non-persistent state duration depend on chemical signal level *ρ*. We use the following equation to fit the mean non-persistent state duration.

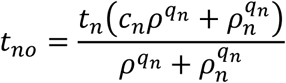

*t_n_* is the non-persistent state duration of cells at low chemical signal area. *q_n_*, *ρ_n_* are parameters to fit the curve. We fit the duration in 1D, using the same signal level used to fit eq. (11).

## Supporting information

**S1 Fig.:**
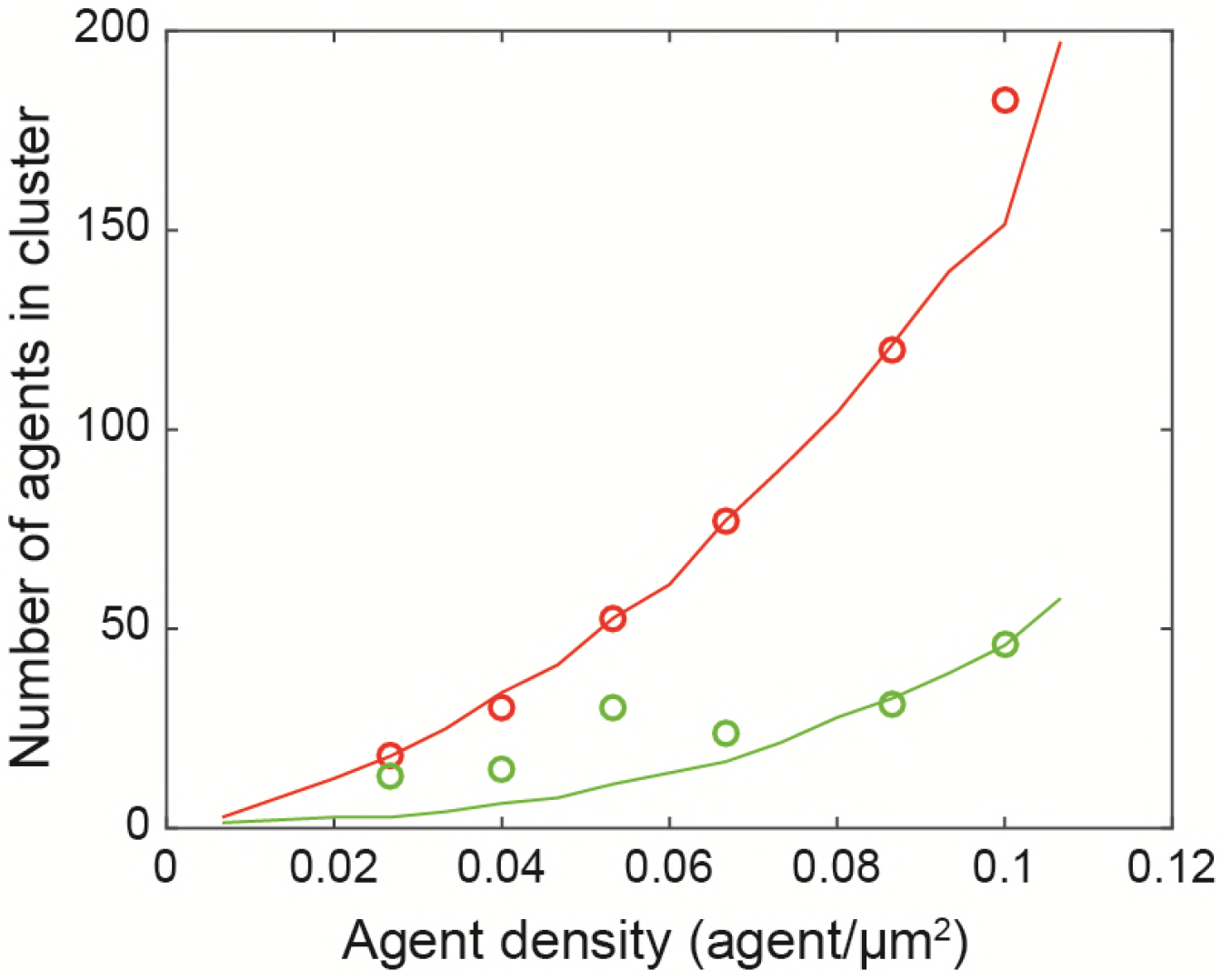
Comparison between our model and the previous Balagam et al. [13] biophysical model. Results of the number of cells in a cluster as a function of cell density for reversing cells. Solid lines are results from Balagam et al. Circles are from our simulation. Red line and markers are for simulation with trail following. Green line and markers are for simulation without trail following.

**S2 Fig.:**
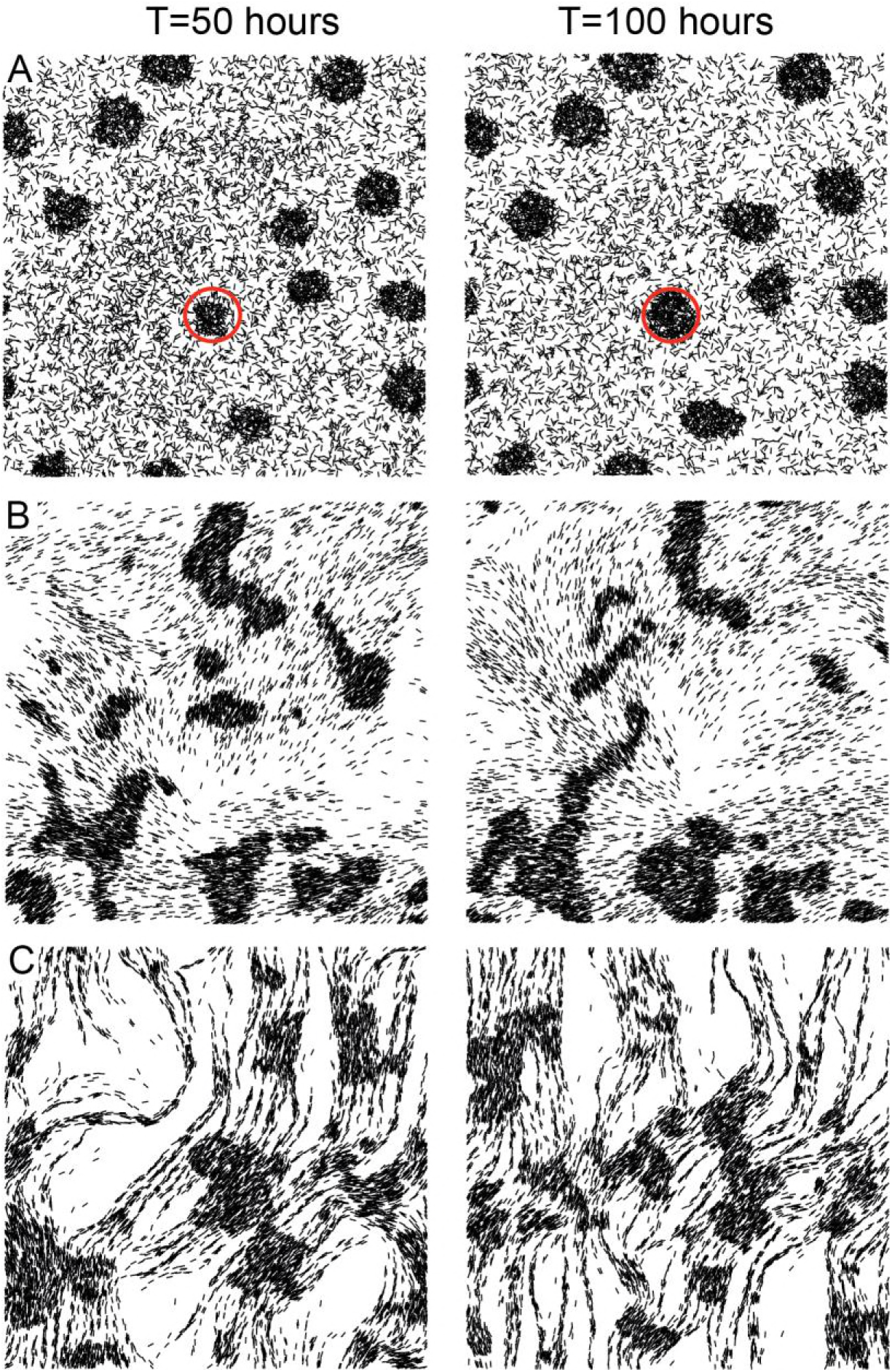
Different aggregation patterns of traffic jam models after long simulations. Left columns: simulations after 50 hours. Right columns: simulation after 100 hours. (A) Agents slow down at high agent density areas. Aggregate in the right circle continues growing between 50 hours and 100 hours. (B) Same as (A), except that agents align moving in a direction with neighbors. (C) Same as (B), except that agents now exhibit trail-following. Agent density is 0.3 agent/μm^2^. Simulation region is 400 μm × 400 μm. For clarity, only 20% of agents are plotted.

**S3 Fig.:**
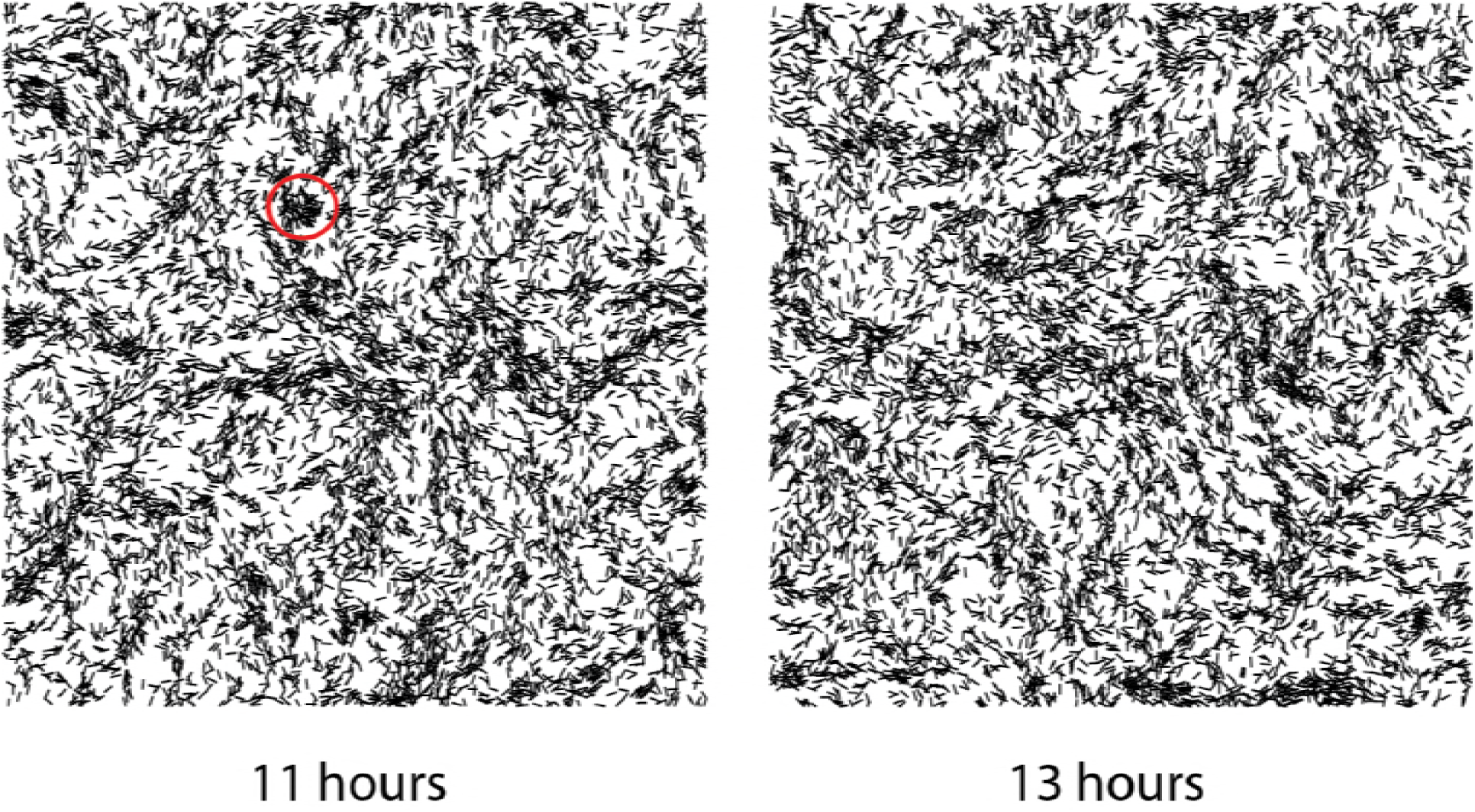
Simulation results of the contact-based signal model at different times. Left panel shows the agents’ pattern at 11 hours, the red circle marks an initial aggregate. Right panel shows agents’ pattern at 13 hours, we observed that the initial aggregate dispersed. Agent density is 0.3 agent/μm^2^. Simulation region is 400 μm × 400 μm. 20% of agents are plotted.

**S4 Fig.:**
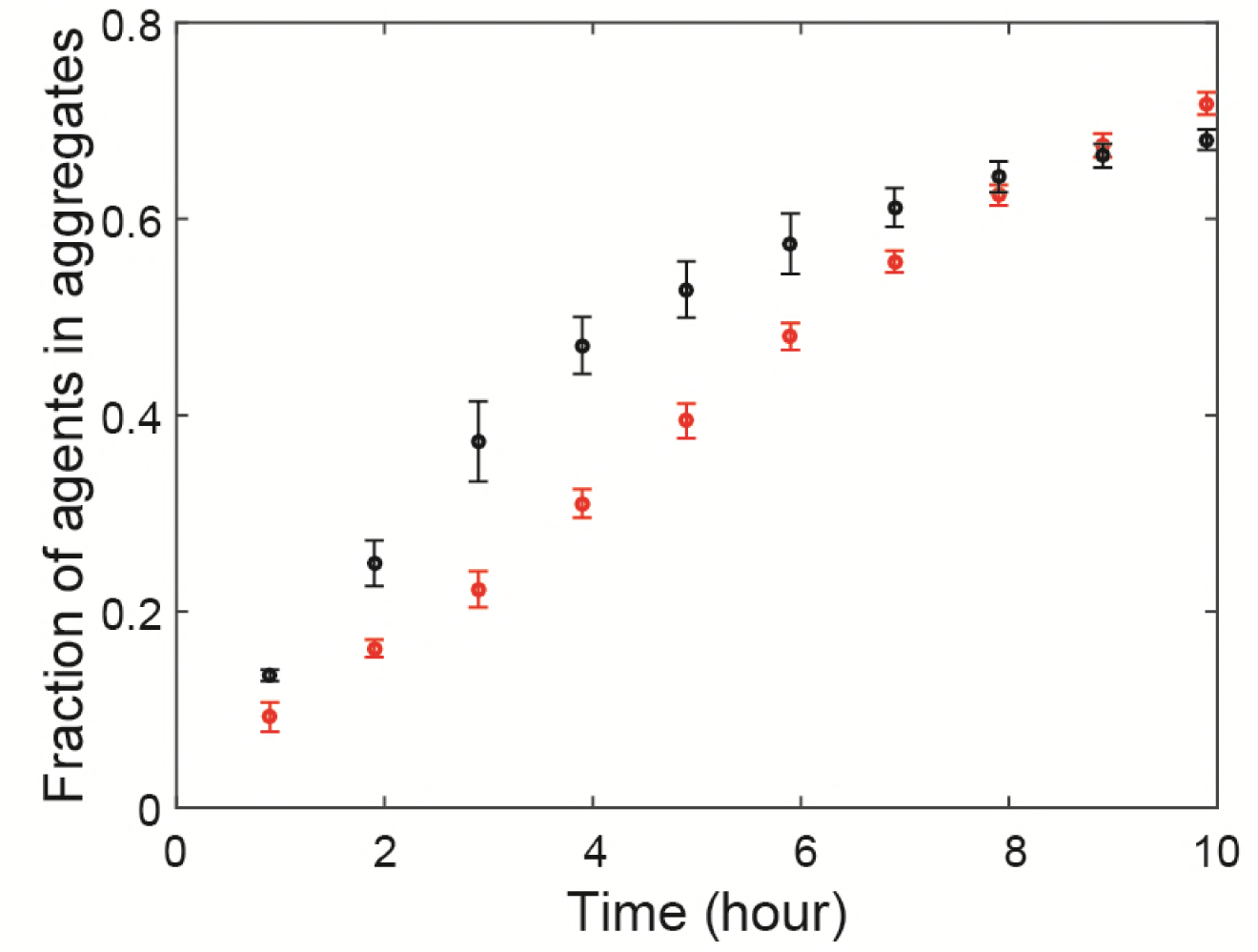
Fraction of agents in aggregate as a function of time in simulation of different agent densities. Agents represent different numbers of cells to keep the overall cell density the same. Black dots are for simulations with 0.3 agent/μm^2^ (each agent represents one cell). Red dots are for simulation with 0.15 agent/μm^2^ (each agent represent two cells). Other simulation parameters are the same as Fig. 5. In our model, agents can represent multiple cells without affecting aggregation speed much.

**S5 Fig.:**
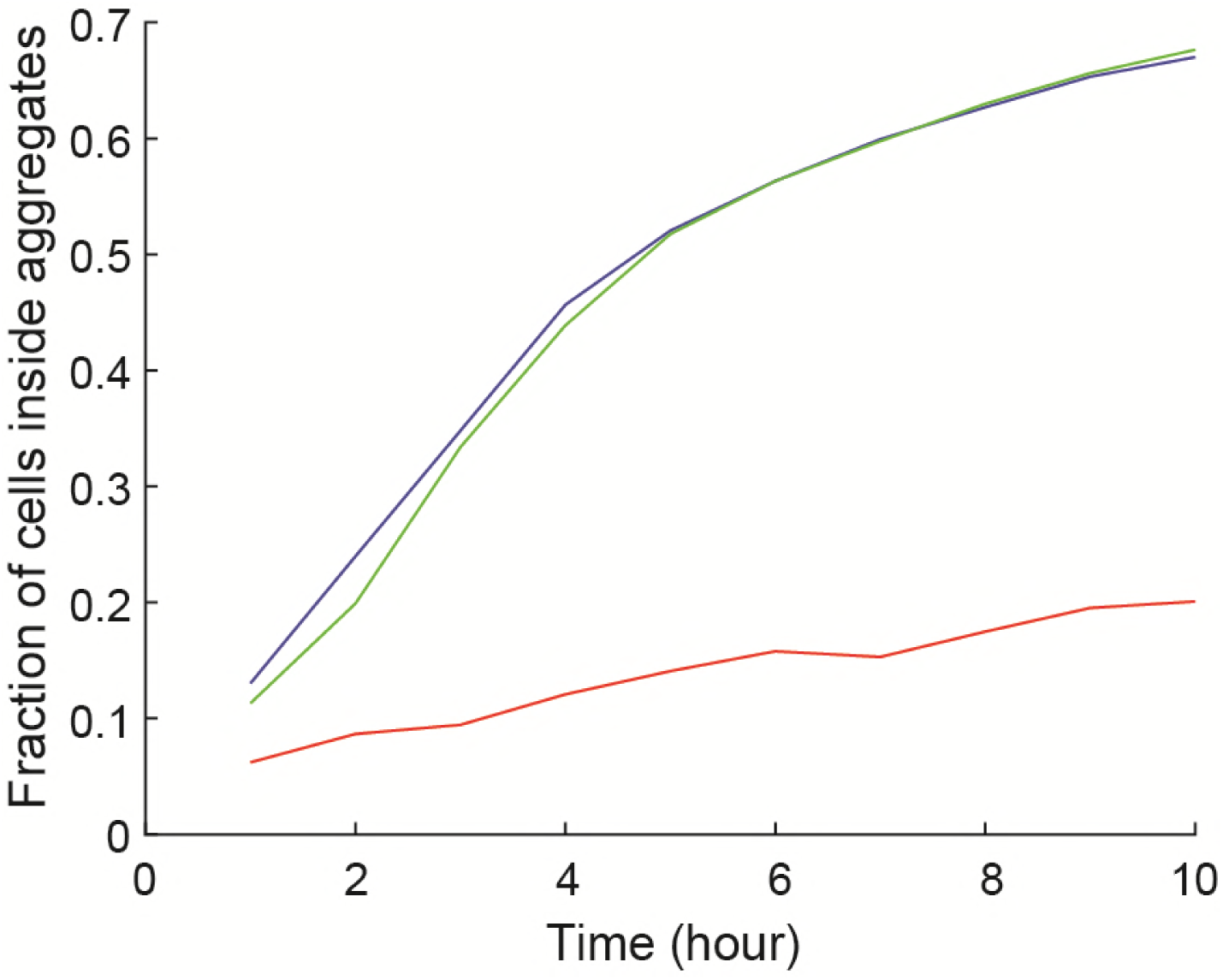
Fraction of cells inside aggregates as a function of time for simulations with different diffusion coefficients. Red line: diffusion coefficient is 3000μm^2^/min. Blue line: diffusion coefficient is 300μm^2^/min. Green line: diffusion coefficient is 30μm^2^/min. Chemical production and decay rate is scaled with diffusion coefficient. Other parameters are same as Fig. 5.

**S6 Fig.:**
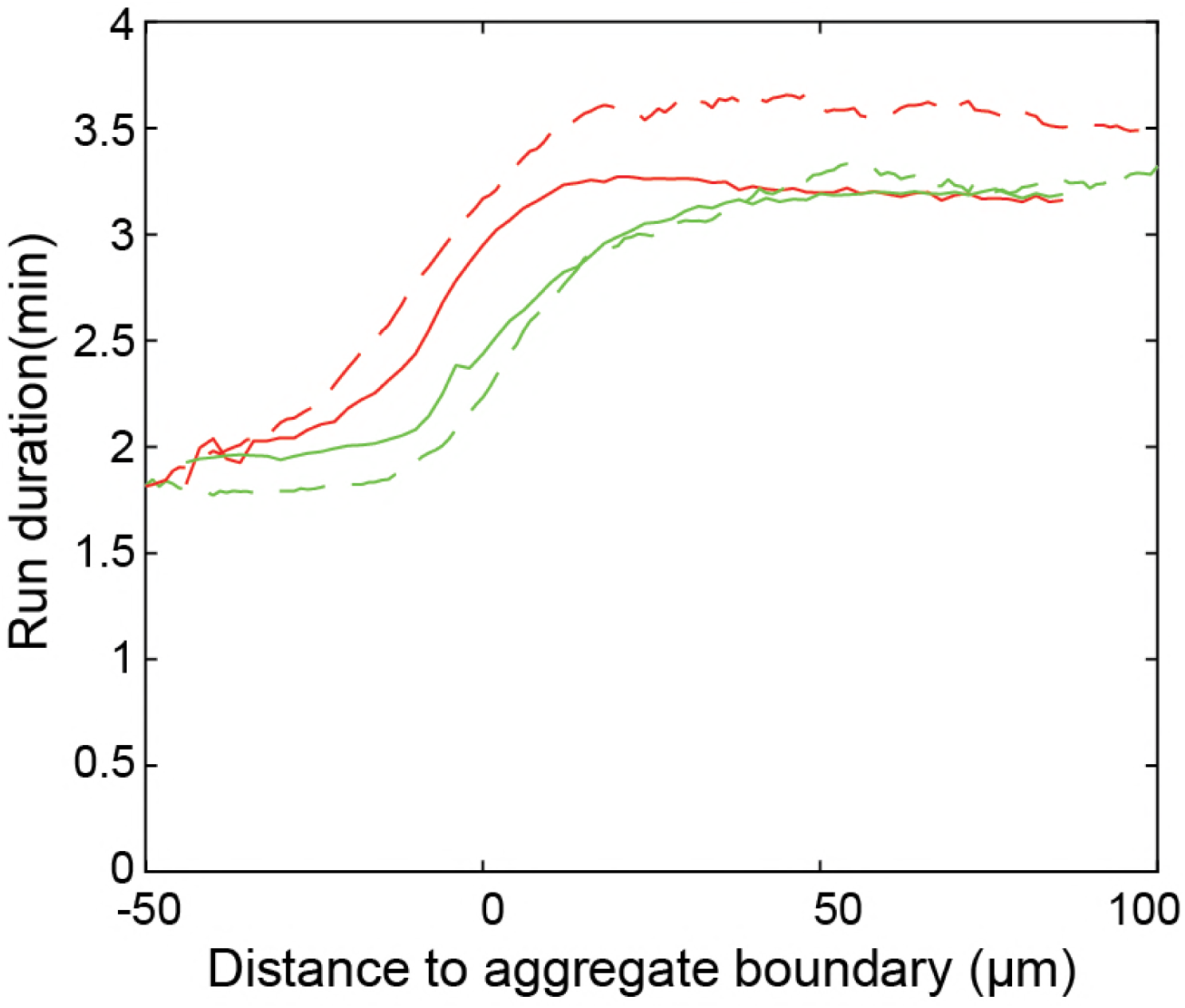
Run duration bias from simulation (solid lines) and experiment (dashed lines). Red lines are runs pointing toward the aggregate centroid, green lines are runs pointed away from the aggregate centroid.

**S7 Table:**
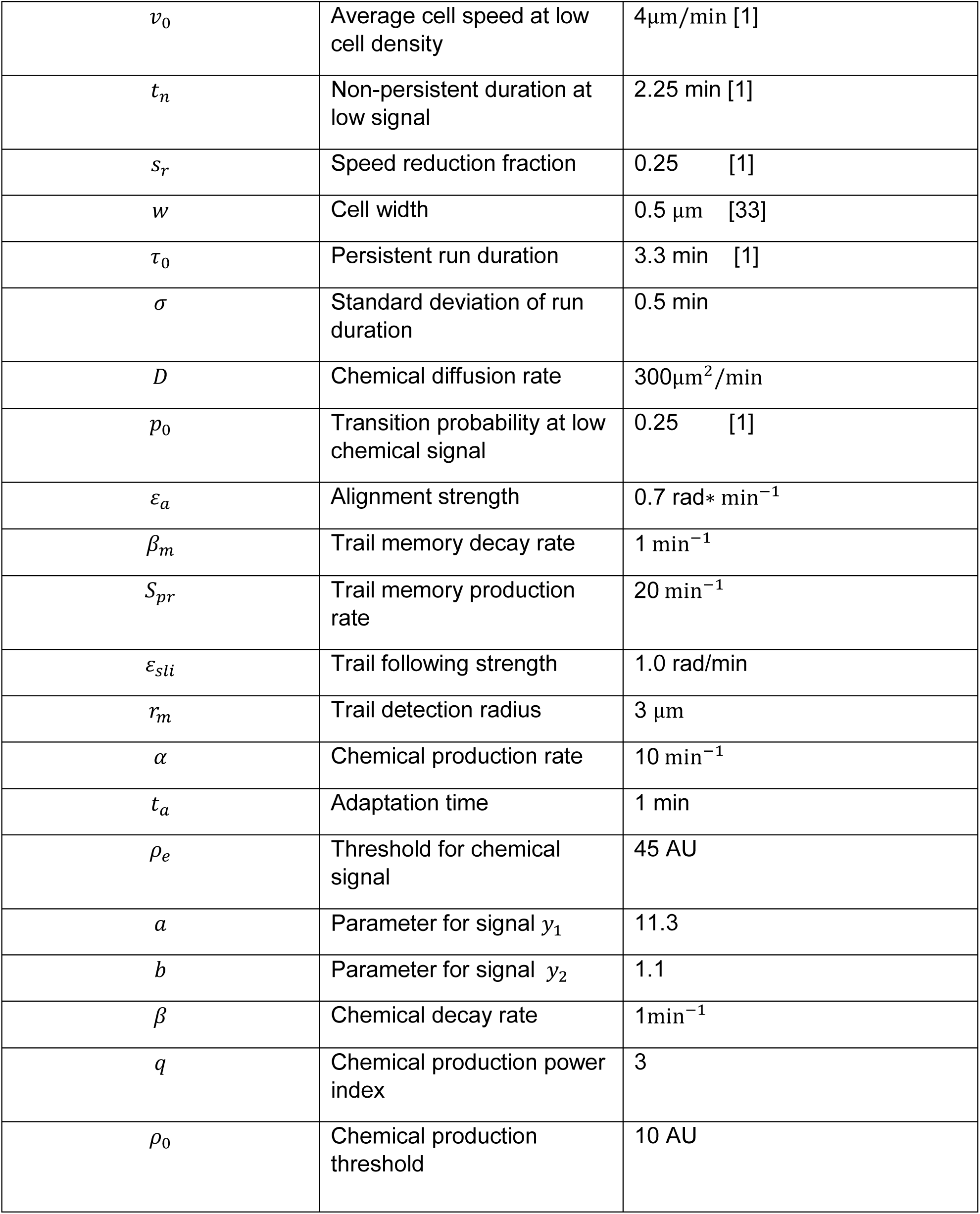
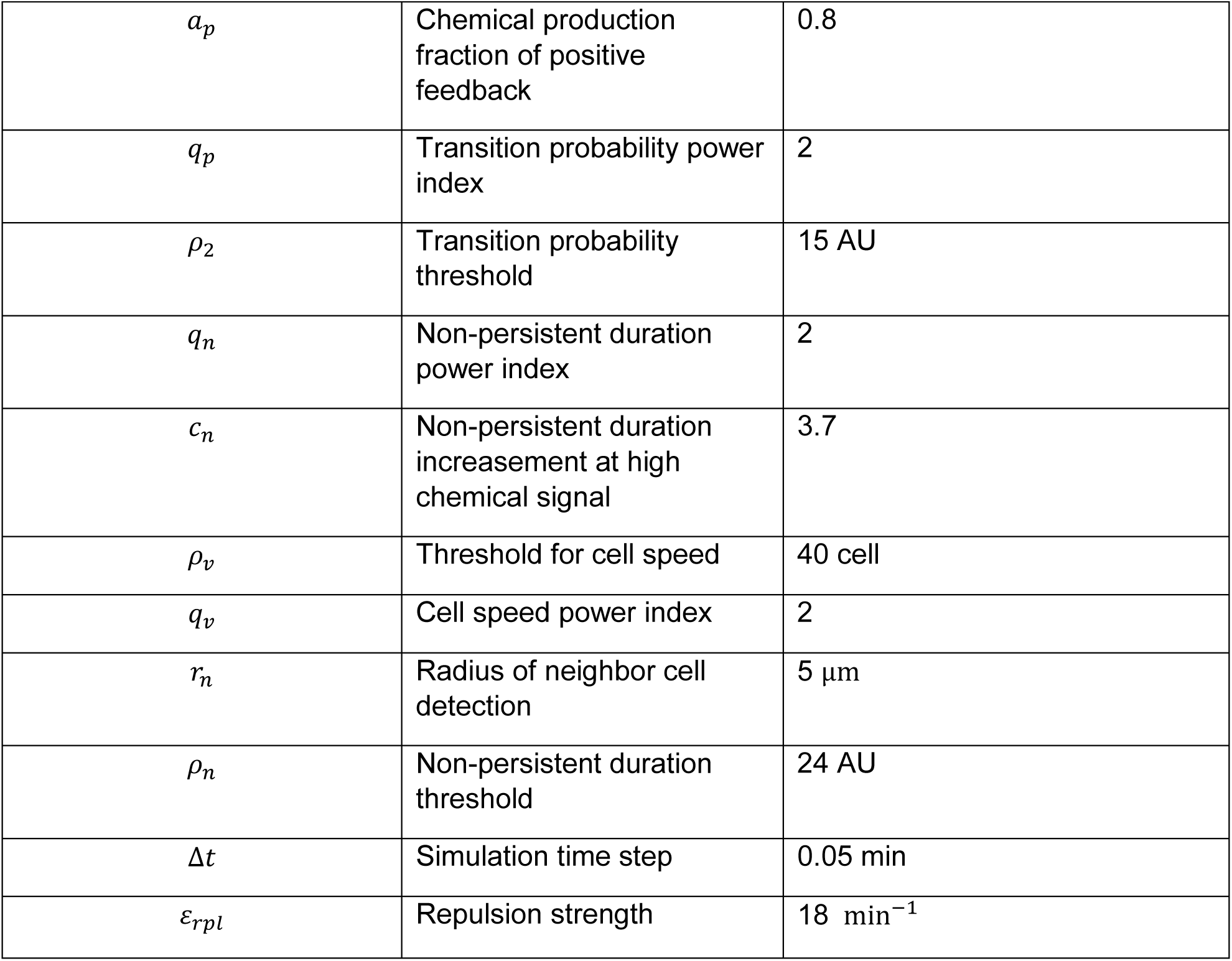
Parameter values used in ABM simulations

